# The Rice XA21 Ectodomain Fused to the Arabidopsis EFR Cytoplasmic Domain Confers Resistance to *Xanthomonas oryzae* pv. *oryzae*

**DOI:** 10.1101/119925

**Authors:** Nicholas Thomas, Nir Oksenberg, Furong Liu, Daniel Caddell, Alina Nalyvayko, Yen Nguyen, Benjamin Schwessinger, Pamela Ronald

**Affiliations:** Department of Plant Pathology and the Genome Center, University of California, Davis, Davis, California, United States of America; The Australian National University, Research School of Biology, Acton ACT 2601, Australia.; Department of Plant Pathology and Microbiology, University of California, Riverside, California 92521, USA

## Abstract

- Rice (*Oryza sativa*) plants expressing the XA21 cell surface receptor kinase are resistant to *Xanthomonas oryzae* pv.*oryzae* (*Xoo*) infection. We previously demonstrated that expressing a chimeric protein containing the EFR (<u>E</u>LONGATION <u>F</u>ACTOR Tu <u>R</u>ECEPTOR) ectodomain and the XA21 endodomain (EFR:XA21) in rice does not confer robust resistance to *Xoo*.
- To test if the XA21 ectodomain is required for *Xoo* resistance, we produced transgenic rice lines expressing a chimeric protein consisting of the XA21 ectodomain and EFR endodomain (XA21:EFR) and inoculated these lines with *Xoo*. We also tested if the XA21:EFR rice plants respond to a synthetic sulfated 21 amino acid derivative (RaxX21-sY) derived from the activator of XA21-mediated immunity, RaxX.
- We found that five independently transformed XA21:EFR rice lines displayed resistance to *Xoo* as measured by lesion length analysis, and showed that five lines express markers of the XA21 defense response (generation of reactive oxygen species and defense response gene expression) after treatment with RaxX21-sY.
- Our results indicate that expression of the XA21:EFR chimeric receptor in rice confers resistance to *Xoo*. These results suggest that the endodomain of the EFR and XA21 immune receptors are interchangeable and the XA21 ectodomain is the key determinant conferring robust resistance to *Xoo*.

## Introduction

Plant cell surface immune receptors confer defense against pathogen infection. Cell surface mediated immunity in plants is mainly conferred by receptor like proteins (RLPs) and receptor like kinases (RLKs) that recognize pathogen associated molecular patterns (PAMPs) (Jones and Dangl 2006; Macho and Zipfel 2014). Three well-studied cell-surface RLKs that confer resistance to bacterial pathogens include FLAGELLIN SENSING2 (FLS2; At5G6330) (Gómez-Gómez and Boller 2000), EF-TU RECEPTOR (EFR; At5g20480) (Zipfel et al. 2006) from *Arabidopsis* and XA21 (U37133) from *Oryza longistaminata* (Song et al. 1995). The identification of the microbial molecules recognized by these three receptors have enhanced in depth characterization of their functional properties. The FLS2 receptor binds the flg22 peptide derived from bacterial Flagellin (Felix et al. 1999; Gómez-Gómez and Boller 2000; Chinchilla et al. 2006). EFR recognizes the elf18 peptide derived from the bacterial Elongation Factor Thermo-unstable protein (EF-Tu) (Kunze et al. 2004; Zipfel et al. 2006). XA21 recognizes the sulfated RaxX (<u>r</u>equired for <u>a</u>ctivation of <u>X</u>a21-mediated immunity <u>X</u>) protein produced by *Xoo* (Pruitt et al. 2015). Although these receptors specifically recognize different molecules, they share similar domain structures including ectodomains containing leucine rich repeats and endodomains containing intracellular kinases of the non-arginine aspartate (non RD) class (Dardick and Ronald, 2006).

Domain swap studies between cell-surface receptors have led to the hypothesis that the nature of the endodomains is the primary determinant dictating the specific disease resistance outcome. For example, studies of a chimeric receptor generated by fusion of the *Arabidopsis* BRASSINOSTEROID-INSENSITIVE1 (BRI1) receptor, which recognizes brassinosteroid hormones, to the XA21 endodomain (BRI1:XA21) (Li and Chory 1997) indicated that the chimeric receptor could be activated by brassinosteroid treatment. Rice cells expressing BRI1:XA21 (NRG-1 in the original publication) and treated with brassinosteroid initiated cell death, produced ROS, and expressed stress-related genes. The stress-related symptoms were attributed to the activation of the XA21 endodomain because the full-length BRI1 receptor does not induce the same stress-related symptoms as BRI1:XA21 (He et al. 2000). These results suggested that the XA21 endodomain was activated upon BRI1 recognition of brassinosteroid and that the specific type of response was most consistent with the response mediated by the XA21 endodomain and not the BRI1 ectodomain.

Another chimera study compared the responses of receptors consisting of ectodomain and endodomain exchanges between EFR and WAK1. WAK1 recognizes oligogalacturonides (OGs) released from damaged plant cell walls (A. Decreux and Messiaen 2005; Annabelle Decreux et al. 2006; Cabrera et al. 2008). Elf18 treated wild-type plants and OG treated plants expressing a WAK1:EFR (WEG) chimeric RLK both produced ROS, ethylene and expressed the EFR-induced genes (At3g22270 and At4g37640) while EFR:WAK1 (EWAK) expressing plants did not. Instead, EWAK plants retained WAK1-like responses by producing ROS but not ethylene in response to OGs (Brutus et al. 2010; Ferrari et al. 2008). WEG and EWAK responses were therefore most consistent with the response conferred by the respective endodomain portion of each fusion protein. Another study showed that fusing the XA21 endodomain to the fungal chitin receptor like protein CEBiP (Kaku et al. 2006) (CRXA-1, and CRXA-3 in the original publication) conferred a more robust immune response to fungal infection by *Magnaporthe oryzae* than when expressing or overexpressing CEBiP alone (Kishimoto et al. 2010). These results suggested that the XA21 endodomain was responsible for conferring the enhanced immune response to *M. oryzae*. Together, these studies indicate that the endodomain of several immune receptors dictate the specific signaling events that lead to disease resistance in whole plants or defense responses in plant cells. These studies also suggest chimeras carrying the XA21 endodomain, when treated with the appropriate ligand, can initiate an immune response similar to that mediated by the full-length receptor XA21.

To further explore the function and specificity of the XA21 endodomain and ectodomain, we previously generated transgenic rice lines expressing EFR, tagged with green fluorescent protein (EFR:GFP), or a chimeric EFR:XA21 protein, consisting of the EFR ectodomain and the XA21 transmembrane and intracellular domain, tagged with GFP (EFR:XA21:GFP) (Schwessinger, Bahar, et al. 2015). Both *EFR:GFP* and *EFR:XA21:GFP* rice plants were susceptible to *Xoo* strain PXO99A and conferred partial resistance to weakly virulent strains (Schwessinger, Bahar, et al. 2015). These studies suggested that although both receptors were capable of recognizing EF-Tu, they were still unable to initiate a robust immune response to PXO99A. As noted in the paper’s discussion, these results were counterintuitive based on earlier domain swap studies that indicated that the endodomain dictates immune signaling and disease resistance (He et al. 2000; Brutus et al. 2010; Kishimoto et al. 2010; Albert et al. 2010).

Although it is unclear why the EFR and EFR:XA21 study conflicted with findings from previous chimeric receptor studies, there are several possibilities to explain these discrepancies. In the case of the EFR:WAK1 and WAK1:EFR study, it could be that the type of kinase domain dictated the distinct signaling mediated by each chimeric receptor because the WAK1 and EFR kinase domains belong to different kinase classes. The WAK1 kinase domain contains an arginine (R) aspartate (D) motif while the EFR kinase domain is non-RD, as described above. The non-RD kinases are almost always associated with immune responses in plants and animals and are likely regulated differently than RD kinases (Christopher Dardick and Ronald 2006; Ronald and Beutler 2010; Chris Dardick, Schwessinger, and Ronald 2012). Thus, the presence of the non-RD domain may dictate an immune response when appropriately activated and the presence of the RD domain may specify a WAK1-like response.

For both BRI1:XA21 and CEBiP:XA21 studies, it is possible that the origin of the kinase domain from XA21 was less important than the fact that the kinase belonged to the non-RD class. For example, it is unclear if fusing BRI1 or CEBiP to other non-RD kinases, such as the kinases from EFR or OsFLS2, would have produced similar results (Takai et al. 2008).

Previous studies have shown that the XA21 ectodomain plays a critical role in the immune response. For example, the *Xa21D* paralog, which lacks a transmembrane and intracellular domain, confer partial resistance to *Xoo* (*Wang et al. 1998*). Unlike *Xa21, Xa21D* only encodes an ectodomain that is nearly identical to the XA21 ectodomain, differing only in 15 amino acid residues compared to the XA21 ectodomain. Similarly, expression of a catalytically inactive variant of XA21, carrying a mutation in the catalytic domain of the kinase (K736E), in rice maintained partial resistance to *Xoo* (Cynthia B. Andaya and Ronald 2003). Together, these studies indicate that the XA21 ectodomain is sufficient to confer partial resistance to *Xoo*, even in the absence of a functional kinase domain.

To further explore the function and importance of the XA21 ectodomain, we generated transgenic rice lines expressing a chimeric protein containing the XA21 ectodomain fused to the EFR transmembrane and intracellular domain, tagged with GFP (XA21:EFR:GFP) (Holton et al. 2015). We found that *XA21:EFR:GFP* rice display robust resistance to *Xoo* strain PXO99A. We also show that *XA21:EFR:GFP* was specifically activated by RaxX as measured by defense response gene expression and ROS production (Pruitt et al. 2015; Schwessinger, Li, et al. 2015; Wei et al. 2016). These results indicate that the XA21 ectodomain and its recognition of RaxX specify robust resistance to *Xoo* even in the absence of the XA21 endodomain.

## Materials and Methods

### Plant material and methods

Rice seeds were germinated on water-soaked filter paper for 5-7 days at 28°C and then transplanted into 2.6-liter pots. Plants were grown in an approximately 80/20 (sand/peat) soil mixture in an environmentally-controlled greenhouse with temperature set between 28-30°C with 75-80% humidity.

### Transgenic rice production

The *Xa21:EFR:GFP* (XA21 aa residues 1-650 fused to EFR aa residues 650-1031) binary vector used in rice transformation was described previously (Holton et al. 2015). Transgenic Kitaake plants expressing the *Xa21:EFR:GFP* transgene were generated by the UC Davis Plant Transformation Facility as described previously (Hiei et al. 1994). pCAMBIA1300 binary vectors carrying the *Xa21:EFR:GFP* construct were transformed into Kitaake calli by Agrobacterium-mediated transformation. Regenerated plants were selected on hygromycin. The presence of the transgene was confirmed in each generation by PCR using transgene specific primers (Table S1).

### Segregation analysis (genotyping, infection, plant conditions)

*Xoo* isolates (PXO99A strain) were plated on peptone sucrose agar plates for 3 days. *Xoo* was suspended in water to approximately 5x10^8^ colony forming units (CFU)/ mL. Greenhouse-grown plants were transported into environmentally controlled growth chambers at the 4 week-old stage. Chamber conditions were set to 26°C, 85% humidity with 12h light/dark cycles. Plants were acclimated to the chamber conditions for 2–3 days before scissor inoculation (Kauffman et al. 1973). The presence of the transgene was identified using PCR genotyping with transgene-specific primers (Table S1).

### Gene expression analysis by qRT-PCR

Total RNA was extracted from detached leaves frozen in liquid nitrogen and powdered using a Qiagen tissuelyser. RNA was extracted from powdered tissue using TRI Reagent and precipitated with isopropanol. RNA was DNase treated using the TURBO DNase kit from Life Technologies. RNA concentrations were normalized to the lowest sample concentration in each experiment. cDNA was synthesized from 2µg of total RNA using the High Capacity cDNA Reverse Transcription Kit by Life Technologies. Gene expression changes were determined by ΔΔCt method (Livak and Schmittgen 2001) normalizing gene expression to Actin (*LOC_Os03g50885*) and using mock treated samples as the reference for stress gene expression. Quantitative real time PCR (qRT-PCR) was performed using a Bio-Rad CFX96 Real-Time System coupled to a C1000 Thermal Cycler (Bio-Rad) using the Bio-Rad SsoFast EvaGreen Supermix. qRT-PCR primer pairs used are described in Table S1. qRT-PCR reactions were run for 40 cycles with annealing and amplification at 62°C for 5 sec and denaturation at 95°C for 5 sec. Single melting curves were observed for all primer pairs used indicating negligible off-target amplification.

### Western Blot Analysis for Protein expression

Anti-GFP (Santa Cruz Biotech) was used to detect EFR:GFP, EFR:XA21:GFP, XA21:GFP and XA21:EFR:GFP. Secondary anti-mouse antibodies (Santa Cruz Biotech) conjugated to horseradish peroxidase were used in combination with chemiluminescence substrates (Thermo) to detect proteins on a Biorad ChemiDoc.

### Reactive oxygen species production

Leaves of 3- to 4-week-old rice plants were cut longitudinally along the mid vein and then into 1 to 1.5 mm thick pieces. Leaf pieces were floated on sterile water overnight. The following morning, two leaf pieces were transferred into one well of a 96-well white plate containing 100 μl elicitation solution (20 μM LO-12 [Wako, Japan], 2 μg/ml HRP [Sigma]). 500 nM of elf18, RaxX21 or RaxX21-sY peptides were used for treatments. ROS production was measured for 0.5s per reading with a high sensitivity plate reader (TriStar, Berthold, Germany).

## Results

### Transgenic rice expressing the XA21:EFR chimeric receptor display robust resistance to *Xoo*

We produced transgenic rice lines expressing an *Xa21:EFR:GFP* chimeric construct to test whether the XA21 ectodomain confers resistance to *Xoo* when fused to the EFR cytoplasmic domain. This construct encodes the XA21 ectodomain (XA21 residues 1-650) fused to the EFR transmembrane, juxtamembrane and cytoplasmic domain (EFR residues 651-1031) with a carboxyl-terminal GFP fusion (Holton et al. 2015) expressed under the maize ubiquitin promoter. We generated 10 independent transgenic T_0_ lines and inoculated the plants with Xoo using a leaf clipping method. We found that 8 of these lines (lines 2, 3, 4, 5, 6, 7, 9, and 10) displayed enhanced resistance to *Xoo* compared with the Kitaake parent line (Fig. S1).

To assess if the resistance phenotype was transmitted to the next generation, we self-pollinated 5 of the 8 T_0_ lines (lines 2,4,5,6,7) and collected T_1_ seed. These T_1_ plants, as well as rice plants expressing and lacking Xa21 as controls, were inoculated with *Xoo* and assessed for resistance by measuring the lengths of disease-induced lesions. We observed that T_1_ individuals that were PCR positive for the transgene in lines 2,4,5, and 6 co-segregated with resistance to *Xoo* (PCR positive to negative ratios 8:4, 21:0, 8:7, and 16:5, respectively). Lesion length averages were approximately 5 cm in resistant individuals compared to approximately 13 cm for susceptible controls (Fig. 1, Fig S2). All T_1_ individuals from line 4 were PCR positive for *Xa21:EFR:GFP* (21:0) which could have been from multiple transgene insertions (*X*^*2*^ (1) = 1.4, p = 0.24) and were resistant to Xoo. All T1 individuals from line 7 were also PCR positive for the *Xa21:EFR:GFP* transgene. However, these plants showed varying degrees of resistance (Fig. S2).

**Figure 1.**
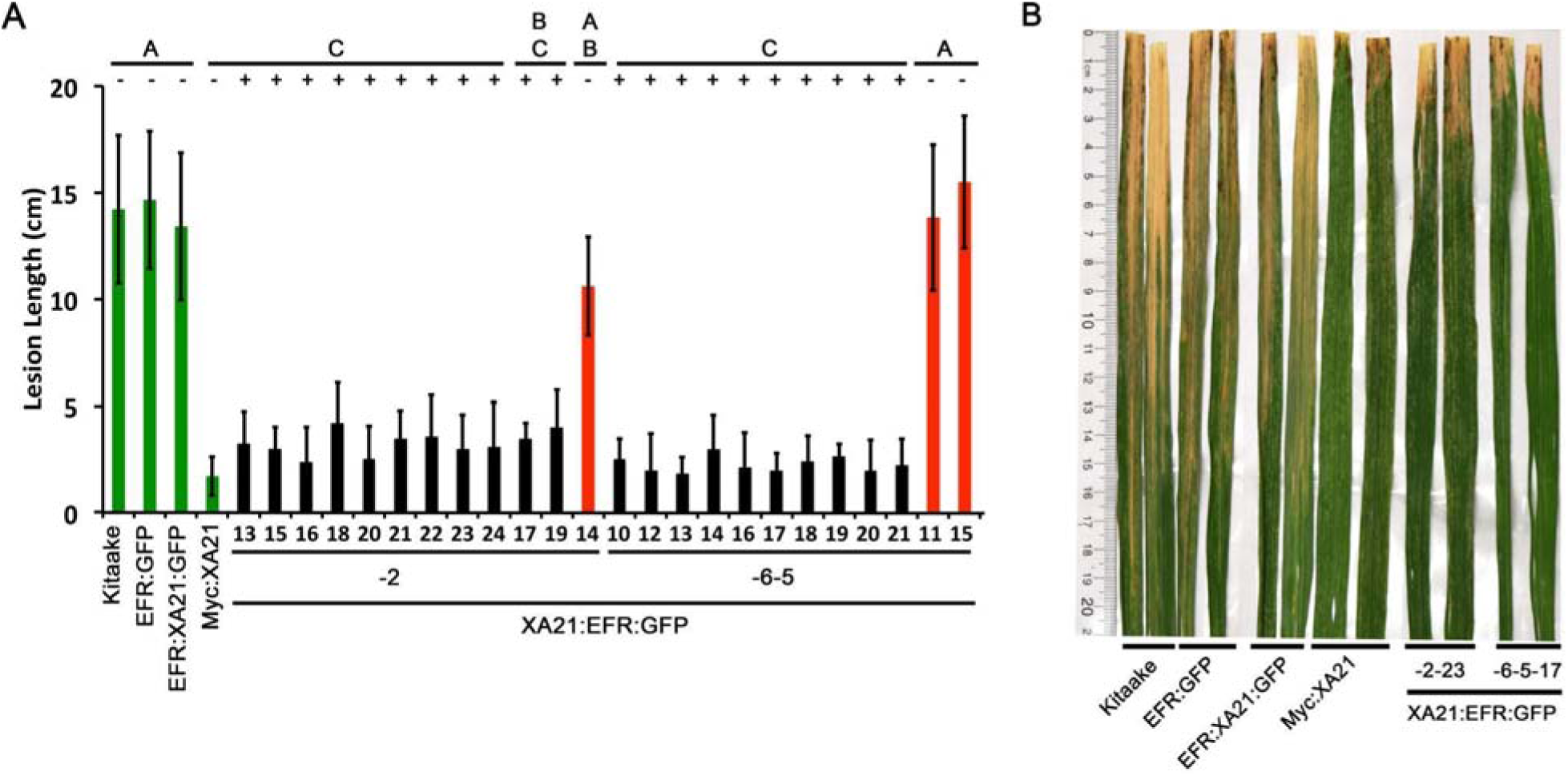
Rice expressing *Xa21:EFR:GFP* are resistant to *Xoo* infection. A, The bar graph represents the average lesion length observed on rice plants infected with *Xoo*. Control lines used were Kitaake, *EFR:GFP, EFR:XA21:GFP*, and *Myc:XA21* rice (green bars). Experimental samples include individuals PCR positive for the *Xa21:EFR:GFP* transgene (black bars) from line 2 and 6 and PCR negative individuals (red bars). Five week old greenhouse grown plants were scissor inoculated with PXO99A (5×10^8^ colony forming units (CFU)/mL) and disease lesions were scored approximately 2 weeks post inoculation. Error bars represent standard deviation from the mean lesion length. Mean lesion lengths are the average of lesion measurements from individual leaves from the same plant (n ≥ 3). Black lines and letters above the graph represent statistical groupings using the Tukey-Kramer HSD test. Different letters indicate significant differences (p < 0.05). This experiment was repeated at least three times with similar results. B, Photograph of select leaves from the same experiment in A. The photograph shows Kitaake, *EFR:GFP, EFR:Xa21:GFP, Myc:Xa21, Xa21:EFR:GFP* individual -2-23, and -6-5-17 leaves infected with *Xoo* and was taken approximately 2 weeks after inoculation.

For subsequent experiments, we focused on two *Xa21:EFR:GFP* lines (-2 and -6) for further molecular characterization experiments (Figs 3, 4, S4). For these experiments, T_1_ plants were used for line 2 and T_2_ plants were used for line 6 (we self-pollinated T_1_ individuals and collected T_2_ seed for line 6) to test if similar phenotypes are observable in different lines and in subsequent generations. We found that T_2_ individuals from line 6 maintained *Xoo* resistance that segregated with the *Xa21:EFR:GFP* transgene (Fig. 1). Because T_1_ and T_2_ individuals from lines 2 and 6, respectively, were still segregating for the transgene, we performed experiments on individual plants that carried the *Xa21:EFR:GFP* transgene, selected by PCR genotyping. We used null segregant individuals as controls.

### The *Xa21:EFR:GFP* chimeric transgene is expressed and XA21:EFR:GFP protein accumulates in stable transgenic lines

We used qRT-PCR to assess if plants containing the *Xa21:EFR:GFP* transgene express the *Xa21* ectodomain and *EFR* cytoplasmic domain. We assessed transcript levels using domain specific primers for regions that encode the XA21 ectodomain, XA21 cytoplasmic domain, and the EFR cytoplasmic domain (Table S1) (Fig. 2A-C). Our results show *Xa21:EFR:GFP*-2-23, -2-24, -6-5-17 and -6-5-18 that carry the transgene specifically express regions encoding the XA21 ectodomain and the EFR cytoplasmic domain. Additionally, these plants do not express regions encoding the XA21 endodomain. Because these plants are not expressing the full-length *Xa21* transcript or endodomain, any immune responses observed in these plants are not mediated by full-length XA21 or the XA21 endodomain.

**Figure 2.**
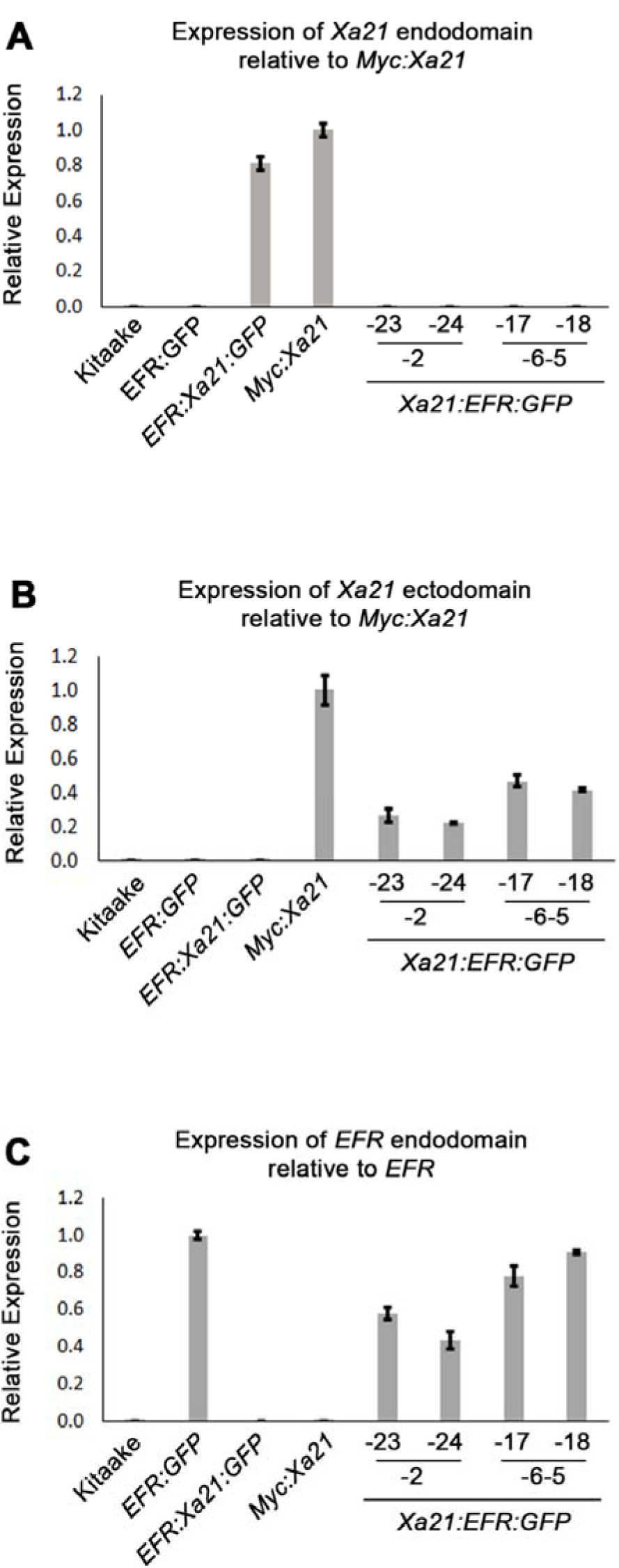
*Xa21:EFR:GFP* transcripts are expressed in stable transgenic lines. Bar graphs represent the relative expression of transgenic transcripts. A, relative amplification of the *Xa21* endodomain with *Myc:Xa21* rice as the expression reference. B, Amplification of the *Xa21* endodomain with *Myc:Xa21* rice as the expression reference. C, Amplification of the *EFR* cytoplasmic domain with *EFR:GFP* rice as the expression reference. Gene expression was measured by quantitative realtime PCR using cDNA amplified from total RNA as a template. Each gene expression measurement is the average of 2 technical replicates and error bars represent the standard deviation between the two measurements.

In addition to the specific *Xa21:EFR:GFP* transcript, we show that XA21:EFR:GFP protein accumulates in transgenic rice. We performed Western Blot analysis to determine if XA21:EFR:GFP protein accumulates in *Xa21:EFR:GFP* transgenic rice using primary anti-GFP antibodies. Our results show that XA21:EFR:GFP protein is detectable in *Xa21:EFR:GFP*-2-28, -2-29, -6-5-4 and -6-5-7 that carry the *Xa21:EFR:GFP* transgene. WT Kitaake and null segregants *Xa21:EFR:GFP-2-32* and *Xa21:EFR:GFP-6-5-6* do not express any GFP tagged protein (Fig. S3). Together, RNA and protein expression indicate that two independent *Xa21:EFR:GFP* transgenic lines express *Xa21:EFR:GFP* transcript and accumulate XA21:EFR:GFP protein.

### RaxX21-sY treated ***Xa21:EFR:GFP*** rice leaves produce reactive oxygen species and highly express stress-related genes

We next assessed if *Xa21:EFR:GFP* rice are able to activate immune responses after RaxX treatments. We used a commercially synthesized, sulfated RaxX peptide, composed of 21 amino acids from the *Xoo* RaxX protein sequence in PXO99A (RaxX21-sY) previously shown to activate XA21-mediated immunity (Pruitt et al. 2015; Wei et al. 2016). Bursts of reactive oxygen species (ROS) are commonly measured to assess immune responses because ROS are rapidly produced as a defense response to pathogen attack (Wojtaszek 1997; Jones and Dangl 2006; Macho and Zipfel 2014). We therefore measured ROS production in *Xa21:EFR:GFP* rice after RaxX21-sY treatment to determine if plants carrying the chimeric protein respond similarly to RaxX21-sY treated plants carrying full-length XA21 (Pruitt et al. 2015). *Xa21:EFR:GFP* rice accumulate ROS in response to RaxX21-sY treatments, but not to mock or elf18 treatments (Figs 3f, S4). In addition, we confirmed that RaxX21-sY treated XA21:GFP rice, expressing the full length XA21 protein tagged with GFP (Fig. S3), accumulate ROS (Fig. 3b). Null segregants did not produce ROS bursts in response to RaxX21-sY treatments (Figs 3e, S4). *EFR:GFP* and *EFR:XA21:GFP* rice responded to elf18, but not to RaxX21-sY, showing that the XA21 ectodomain in full-length XA21 and XA21:EFR:GFP proteins is necessary for RaxX-triggered immune responses (Figs 3c and 3d).

**Figure 3.**
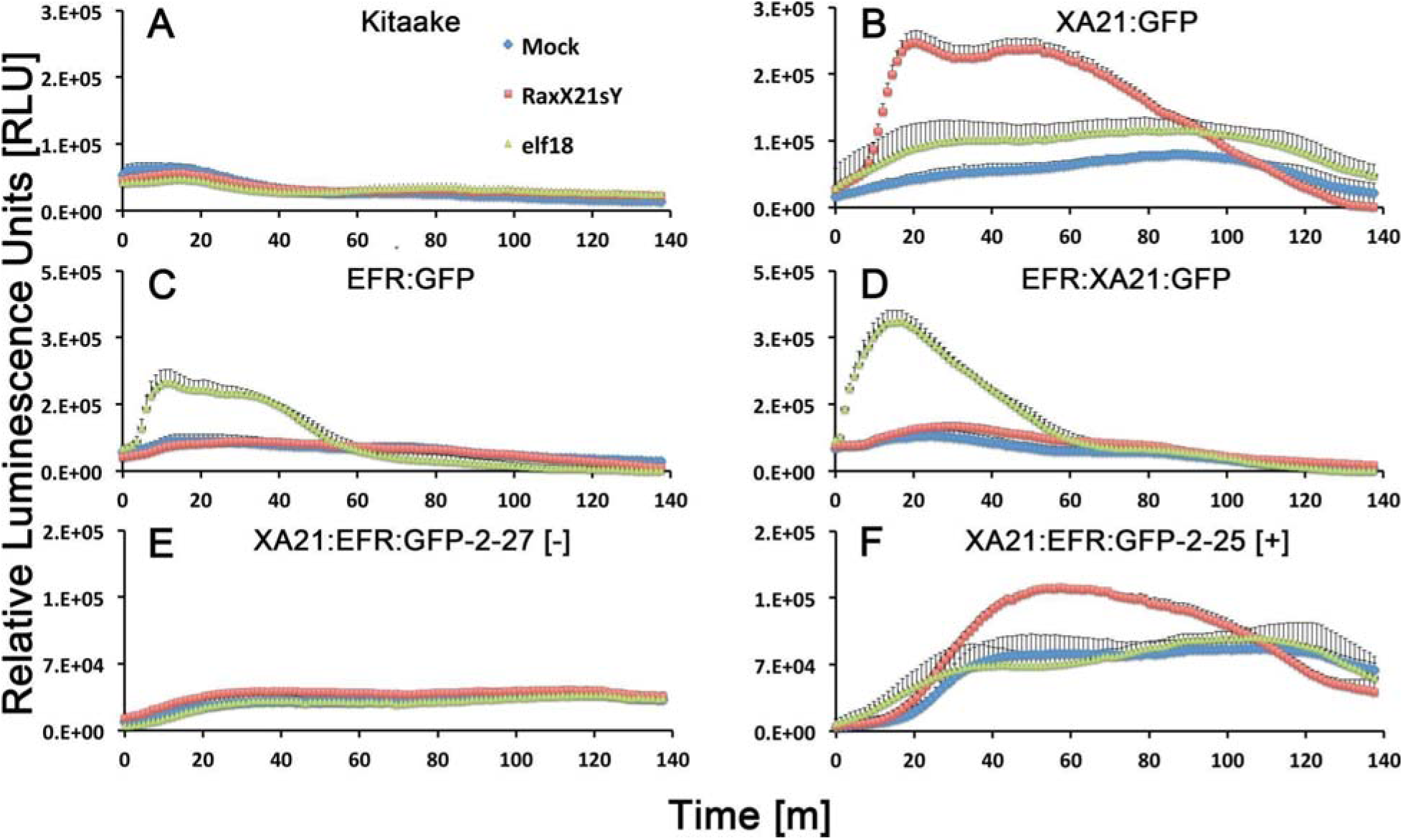
Reactive oxygen species accumulate after peptide treatments. Reactive oxygen species (ROS) production after water (mock, blue diamonds), 500 nM RaxX21-sY (red squares), or 500 nM elf18 peptide treatments (green triangles). A, ROS production in wild-type Kitaake rice and B, *Xa21:GFP* rice C, *EFR:GFP* rice and D, *EFR:Xa21:GFP* rice. E, ROS production in a T_1_ null-segregant individual from *Xa21:EFR:GFP* line -2. F, ROS production in a T_1_ individual from line -2 that segregates for the *Xa21:EFR:GFP* transgene. Each datapoint represents an average of four technical replicate measurements and error bars represent the standard error of the averages. These experiments have been repeated three times with similar results.

We next measured stress-related marker gene expression in RaxX21-sY treated *Xa21:EFR:GFP* rice to further characterize the XA21:EFR:GFP-mediated response. We measured the expression of rice defense marker genes *PR10b, LOC_Os02g36190, LOC_Os06g37224*, and *LOC_Os11g42200* (Chen et al. 2014; Pruitt et al. 2015; Thomas et al. 2016) using a detached leaf treatment assay. RNA was extracted from detached leaves of 4 week old plants mock treated with water or with 500 nM of RaxX21-sY for 6 hours. Gene expression was measured in individuals XA21:EFR:GFP-2-23 and -6-5-17 by quantitative real-time PCR. Higher expression was observed in each of the stress-related genetic markers in RaxX21-sY treated *Myc:Xa21* and *Xa21:EFR:GFP* rice (Fig. 4a-d). Gene expression was not induced in any mock treated samples or Kitaake samples treated with RaxX21-sY. Individual 2-23 only showed higher induction of *LOC_Os11g42200* and *LOC_Os06g37224*, and only induction of *LOC_Os06g37224* was significant (Fig. 4c). Together, the results from ROS and gene expression experiments suggest that the XA21 ectodomain in XA21:EFR:GFP is sufficient to recognize RaxX and that the EFR endodomain can be substituted for the XA21 endodomain to transduce immune responses after RaxX treatment.

**Figure 4.**
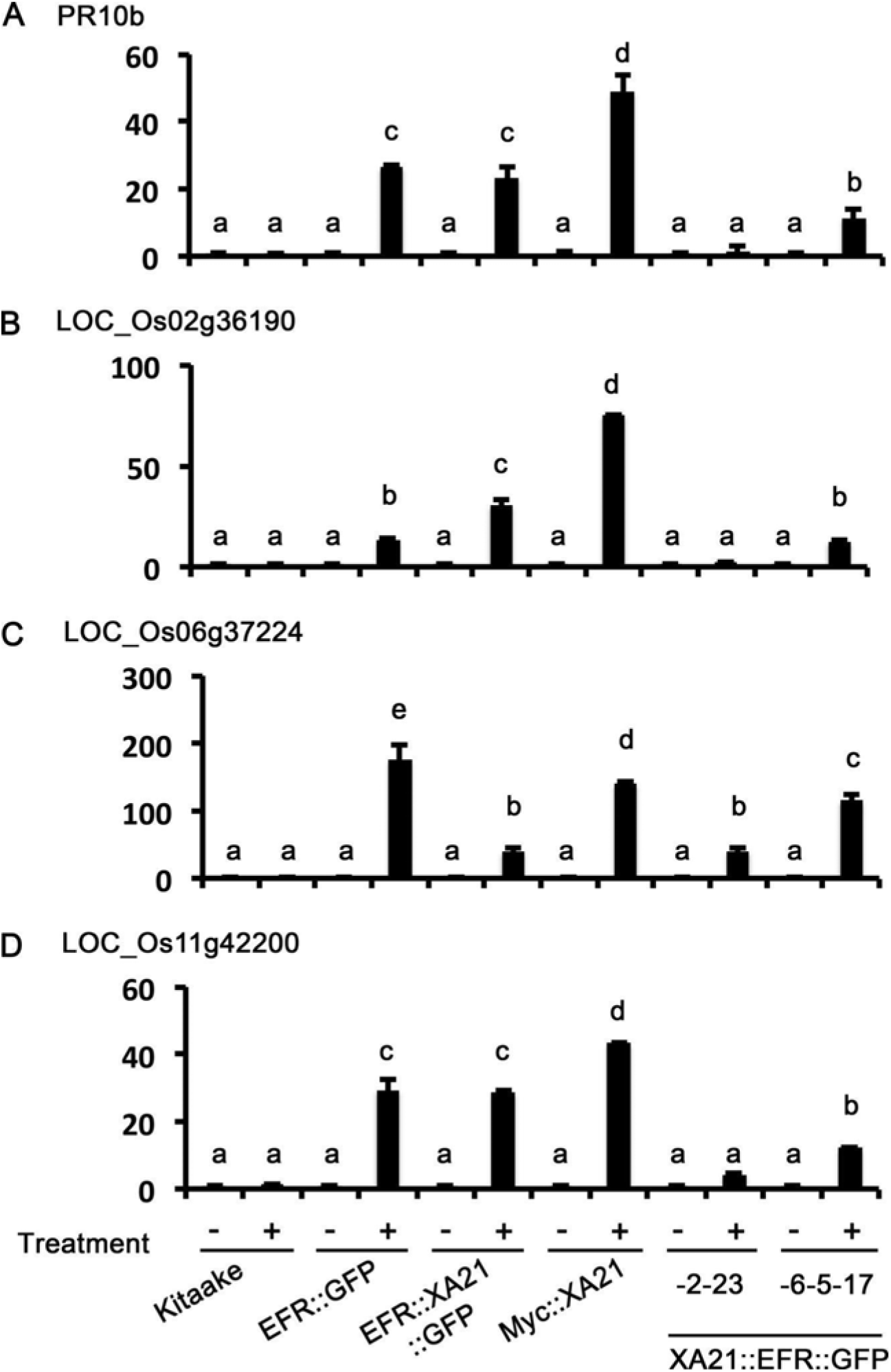
*Xa21:EFR:GFP* rice express stress related genes after RaxX21-sY treatment. Gene expression profiles of four stress-related genes, *PR10b, LOC_Os2g36190, LOC_06g37224*, and *LOC_Os11g42200*. Samples are rice leaves from wild-type Kitaake, Myc:XA21 rice, and individuals -23 from *Xa21:EFR:GFP* line -2 and individual -17 from line -6-5. Leaves were mock treated with water (-) or with 500 nM RaxX21-sY (+). Letters indicate significant difference in gene expression compared to mock using the Tukey-Kramer HSD test (alpha = 0.05). Expression levels are normalized to mock treatment of the same line. Bars depict average expression level relative to actin expression ± standard error of three technical replicates. This experiment was repeated twice with similar results.

## Discussion

Here we show that the ectodomain of XA21 is sufficient to confer full resistance to *Xoo* strain PXO99A when fused to the intracellular domain of the Arabidopsis immune receptor EFR (Figs 1, S1, S2). We previously demonstrated that a functional EFR:XA21:GFP is not able to confer resistance to *Xoo* when expressed in rice. Together these results suggest that the XA21 extracellular domain and the recognition of RaxX are the key properties that dictate the robust immune response of XA21. Both the native XA21 endodomain as well as the EFR endodomain fused to the XA21 ectodomain appear to be interchangeable as both XA21 and EFR kinases can confer robust resistance when fused with the XA21 ectodomain. This result slightly contrasts with previous domain swap studies that indicated that the endodomains of immune receptors were the defining properties of the immune receptor responses (He et al. 2000; Brutus et al. 2010; Kishimoto et al. 2010; Albert et al. 2010). Although XA21 mutants and XA21 derivatives that lack a functional kinase domain maintain partial resistance, it appears that a functional kinase domain is required for robust resistance (C. B. Andaya and Ronald 2003) (Wang et al. 1996).

Despite the evidence that rice expressing XA21:EFR:GFP are resistant to Xoo, it is unclear why plants expressing the reciprocal EFR:XA21:GFP protein are susceptible to *Xoo* (*Schwessinger, Bahar, et al. 2015*). We hypothesize that the XA21 ectodomain is critical for conferring robust resistance because it interacts with additional rice specific signaling components that the EFR ectodomain is unable to bind. In partial support of this hypothesis, we previously showed that the EFR kinase domain does not interact with some of the previously identified XA21 kinase domain signaling components, including the negative regulator XB15 and positive regulator XB3 (Schwessinger, Bahar, et al. 2015). Future studies might be aimed at identifying these elusive ectodomain specific signaling partners to better understand XA21-mediated immunity.

## Acknowledgments

We would like to thank Dr. Nicholas Holton and Prof. Dr. Cyril Zipfel from the Sainsbury Laboratory for providing the *Xa21:EFR:GFP* construct.

## Funding

This project was funded through NIH grant GM59962 and the NSF PGRP grant IOS-1237975. B. S. was supported by a Human Frontiers Science Program long-term postdoctoral fellowship (LT000674/2012) and a Discovery Early Career Research Award (DE150101897).

